# Context-dependent Rhythmicity in Chimpanzee Displays

**DOI:** 10.1101/2024.10.10.617583

**Authors:** Bas van der Vleuten, Veerle Hovenkamp, Judith Varkevisser, Michelle Spierings

## Abstract

Rhythm is an important component of human language and music production. Rhythms like isochrony (intervals spaced equally in time), are also present in vocalisations of certain non-human species, including several birds and mammals. This study aimed to identify rhythmic patterns with music-based methods within display behaviour of chimpanzees (*Pan troglodytes*), humans’ closest living relatives. Behavioural observations were conducted on individuals from two zoo-housed colonies. We found isochronous rhythms in vocal (e.g. pants, grunts and hoots), as well as in motoric (e.g. swaying and stomping) behavioural sequences. Among individuals, variation was found in the duration between onsets of behavioural elements, resulting in individual-specific tempi. Despite this variation in individual tempi, display sequences were consistently structured with stable, isochronous rhythms. Overall, directed displays, targeted at specific individuals, were less isochronous than undirected displays. The presence of rhythmic patterns across two independent colonies of chimpanzees, suggests that underlying mechanisms for rhythm production may be shared between humans and non-human primates. This shared mechanism indicates that the cognitive requirements for rhythm production potentially preceded human music and language evolution.

## Introduction

Rhythm plays a key role in the production and perception of language and music in humans. The foundation of most rhythms present in language and music, is isochrony: every consecutive interval between actions, has approximately the same duration [1]. This form of consistency even appears in basal physiological processes, such as a beating heart and respiration [2,3]. In the last few decades, a stronger focus on the comparative approach has led to increasingly more non-human animals being studied for their rhythmic capacities. Research in the field of comparative behavioural studies has shown that traits for perception and production of simple (e.g. isochronous) rhythms seem to be universally present in a broad range of species [4–6]. More elaborate rhythmic capabilities, for instance entrainment to an external sound pulse, are likely exclusive to humans and relatively few other species [7–10]. This universal base for rhythm perception and production, with exception of advanced skills in a few species, points towards a gradual process of small cognitive and motoric changes over time that eventually evolved into the elaborate rhythmic capacities used in language and music [11]. However, to understand the cognitive underpinnings of rhythmicity and to form evolutionary theories of rhythmicity, additional empirical data from a comparative perspective is necessary.

Rhythm seems to permeate behaviours across species, nevertheless, there is a difference between rhythm present in intentional, goal-oriented behaviours, rooted in deeper cognitive processes, and behaviours that are not actively controlled (e.g. an isochronous heartbeat). To quantify rhythm in intentional animal behaviour, a generalised music-based method was used in a study by Roeske and colleagues [12], who researched the similarities between birdsong and human music. Similarly to human music, the song of thrush nightingales and zebra finches has an underlying, isochronous structure, with additional categorical rhythms, such as small-integer ratios of 1:2, where consequent intervals have twice the duration of prior intervals. Likewise, using this or a highly similar method, spontaneously-produced, rhythmic vocalisations have been quantified in a few mammals: bats (*Saccopteryx bilineata*) [13], seal pups (*Phoca vitulina*) [14], hyraxes (*Procavia capensis*) [15], lemurs (*Indri indri)* [16], gibbons (*Hylobates lar* [17]; *Nomascus gabriellae, Nomascus leucogenys* and *Nomascus siki* [18]) and orangutans (*Pongo pygmaeus wurmbii*) [19]. These primates (lemurs, gibbons and orangutans) show isochrony or other categorical rhythms in their song, further indicating that the evolutionary origins of these traits emerged before the lineage split between humans and other primates. Moreover, recent efforts to study isochrony in motoric behaviour, rather than vocalisations, found a significant tendency towards isochrony in the grass-plucking movements of wild geladas [20]. These promising results show that these methods are suitable for quantifying rhythmic behaviour. Similar approaches will help gather more data, enabling comparisons between species, advancing the knowledge of the evolutionary trajectory of rhythm production.

Currently, rhythm production and perception still need to be assessed across the majority of the primate species, although earlier research has suggested temporal consistency within primate social behaviour. Regarding humans’ closest living relatives, chimpanzees (*Pan troglodytes*) drum on buttresses of trees with steady cadence in the wild [21]. This behaviour is commonly used for locating and coordinating subgroups over longer distances, during moving events [21], with individual- and group-specific variation to ensure clear recognition within and between groups [22]. The chimpanzees select buttressing tree roots with specific sound propagating features, indicating that it is a non-random, intentional behaviour [23]. These drum sessions are often accompanied by pant-hoot call sequences, which have similar inter-individual and group-specific variation as the buttress drumming behaviour [22,24,25]. Besides regularity in long-distance communication, there has been a recorded instance of a zoo-housed chimpanzee drumming on a barrel within its enclosure, in a constant pattern with occasional changes in tempo [26]. Even though observations as described in this anecdotal report have not frequently been reported ever since, it does further enhance the likelihood of rhythmic patterns, similar to those found in language and music, being present in chimpanzee behaviour.

This study focuses on rhythm in the structured display behaviour of zoo-housed chimpanzees. In display behaviour, (mostly) male chimpanzees use an elaborate repertoire of vocalisations and motoric movements [27,28], to establish dominance, intimidate hierarchical challengers, resolve conflict, attract a mate or play [27,29–31]. The sequences of recurring movements, such as drumming, chest beating and swaying, are accompanied by frequent vocalisations, like pants, hoots, grunts and screams. These multimodally produced behavioural sequences form the ideal opportunity for studying rhythm in these great apes. Display behaviour can be either directed, with optical and bodily orientation towards the target individual(s) [29], or undirected, with no immediate target, which is rather a show of strength, endurance or frustration towards the entire group [32]. When rhythm can be deducted from display behaviour sequences, it is also possible to assess whether this rhythmicity remains consistent or varies across different contexts.

We measured rhythmicity in chimpanzee display behaviour, by extracting the inter-onset-intervals (the intervals between the starts of two consecutive vocalisations or movements) and ratios between different behavioural elements, a well-established method for rhythmic analyses [12,33]. We have taken into consideration whether these potential rhythmic structures were actively produced or the results of physical limitations, by considering inter-individual patterns and tempo variations between the displays of each individual. Besides that, other factors that might influence rhythm production, such as the mode of production (vocal or motoric) and individual variation, were examined. Quantifying rhythm and understanding which factors influence rhythm in chimpanzee behaviour, provides a greater understanding of whether certain aspects of rhythm perception and production are universal across species. Adopting this generalised method in comparative rhythm research, allows us to form evolutionary theories of rhythmicity, which has been rudimentary to the evolution of human language and music.

## Material and methods

### Animals and recordings

The data were collected by observing the chimpanzees (*Pan troglodytes*) of two colonies housed in Beekse Bergen (Hilvarenbeek, The Netherlands) and Burgers’ Zoo (Arnhem, The Netherlands). In total 43 individuals were observed, 26 individuals resided in Beekse Bergen and 17 individuals in Burgers’ Zoo. Though the age-ranges of the two colonies are similar, the individuals housed in Burgers’ Zoo were on average twelve years older. Furthermore, a substantial number of individuals, housed in Beekse Bergen, have been part of a control group for medical research at the Biomedical Primate Research Centre (BPRC, Rijswijk, The Netherlands). All other observed individuals were born and raised in their current, or other, zoos.

Each colony was divided into two separate groups which were within audible distance of each other. The groups occupied enclosures with in- and outside habitats, however, observations were only done outside to accurately record the displays. All behaviours were recorded using JVC Everio GZ-R415 video cameras, a Sennheiser directional microphone and a Marantz Professional PMD661 audio recorder, during all-occurrence sampling sessions that lasted up to five hours. Lengths of sessions depended on the time spent outside by the individuals. In total nearly 90 hours of recordings were collected, spread over 35 days (April-May in Beekse Bergen, September-November in Burgers’ Zoo). Only vocalizations where the individual that emitted the vocalisation could be clearly identified were included in the study. This resulted in 132 displays (included: *vocal* = 73, *motoric* = 110) performed by 29 individuals. Additional information regarding group composition and individuals’ contribution of displays, can be found in the Supplementary Material, S1,2.

### Visual and acoustic analysis

Before analysing, raw videos and audio recordings were synchronised and individual display sequences were then isolated from the continuous recordings, with video-editing software [34,35]. The isolated WAV audio files were annotated with Praat 6.3.10 [36] (*vocal*) and MP4 video files with BORIS 7.13.6 [37] (*motoric*). The onset and offset of all calls (by call types: grunts, hoots, screams, barks, pants) or motoric actions (see Ethogram - Supplementary Material, S3) were noted within the display sequences. The onset-times were then used to calculate the inter-onset-intervals (t_k_) between consecutive behavioural elements (i.e. vocalisations and motoric movements). To assess the variability of these intervals, the coefficient of variation (ratio of the standard deviation to the mean of the inter-onset-intervals in a sequence) was computed for each behavioural sequence. Because of the small number of intervals in some of the sequences, we used the formula for an unbiased estimator [33,38]. A lower coefficient of variation indicates more regular intervals in a sequence. To quantify rhythmic patterns within the behavioural sequences, the ratio (r_k_) between consecutive inter-onset-intervals was computed by 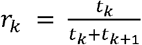. Here, the calculated interval-ratios translate to numeric values between 0 and 1, where an interval-ratio of 1:1, r_k_ = 0.500, marks isochrony. Besides isochrony, other categorical rhythms with ratios of 1:3 (r_k_ = 0.250), 1:2 (r_k_ = 0.333), 2:1 (r_k_ = 0.667) and 3:1 (r_k_ = 0.750), known as small-integer-ratios (SIR), were analysed. Interval-ratios were considered integer (rhythmic) when they fell within the pre-set boundaries of the categorical rhythms [12]. Off-integer (arrhythmic) interval-ratios, grouped outside the rhythmic ranges, form the counterparts of the integer interval-ratios.

### Statistical analysis

All statistical analyses were performed in R 4.2.2 [39]. To quantify rhythmicity in the display behaviour, we adopted the methodology previously proposed by Roeske and colleagues [12], Anichini and colleagues [14] and De Gregorio and colleagues [16]. First, the calculated interval-ratios between consecutive elements were grouped into bins of the categorical rhythms. Here, only individuals with at least 50 interval-ratios across the entire dataset were included to make a proper comparison. The modes of production (vocal/motoric) were also separately analysed, including only individuals with at least 25 interval-ratios in the corresponding mode (Supplementary Material, S1,2). Following this, the interval-ratios of the pre-selected individuals were visualised in density plots (Data visualisation; R packages *ggplot2* [40], *gridExtra* [41], *reshape* [42], *patchwork* [43]). To assess whether the observed distribution could be a product of chance, we compared it with a generated uniform distribution of interval ratios based on the minimum and maximum values of the observed behaviour, following Anichini and colleagues [14] and De Gregorio and colleagues [16]. This simulates a null ratio distribution without rhythmic categories. For this comparison an asymptotic two-sample Kolmogorov-Smirnov test was used to determine the likelihood of the observed interval-ratios originating from the referential uniform distribution [14].

Furthermore, for each individual, counts of interval-ratios within integer and off-integer categories were divided by the categorical bin size, to normalise for the variable bin sizes between categories. Then, the density of each categorical integer rhythm (i.e. isochrony or small-integer-ratios) was compared with their neighbouring off-integer counterparts, using paired Wilcoxon signed-rank tests.

Next, the average interval-duration per display for each individual was calculated and plotted, to visualise the inter-individual variation of temporal properties in display-sequences. For this comparison between individuals, only those with at least four performed displays were selected from the previously selected group. This prevents biased outcomes where the appearance of high consistency in individuals with fewer displays, is merely the result of having less intervals.

The influence of zoo, mode of production and directedness of the display on the regularity of the inter-onset intervals was assessed with a linear mixed model (LMM; R package *lme4* [44]; function *lmer* [45]). The response variable in this model was the unbiased coefficient of variation of the different behavioural sequences, and the fixed effects were the housing-zoo of the focal individuals (Beekse Bergen or Burgers’ Zoo), the mode of production (i. e. vocal or motoric displays), and directedness of the display (directed or undirected). The displaying individual was added as a random effect factor.

Moreover, the influence of these factors on the observed rhythmic patterns was assessed with a generalised linear mixed model (GLMM; R package *lme4* [44]; function *glmer* [45]). In this model, the response variable was the individual counts of isochronous interval-ratios, or isochrony rate, which was related to the model via a binomial link function (1 = isochronous ratio, 0 = non-isochronous ratio). The model incorporated the corresponding colony of the focal individuals, mode of production and directedness of the display as fixed effects. Random effect factors were the displaying individual and the specific display sequence (i.e. individual-ID and display-ID). Other factors, such as demography (age, sex), intergroup variation (within zoo colony) and behaviour type (call types/ motoric behaviour categories), did not improve the fit of the model (Model assessment; R packages *performance* [46], *DHARMa* [47]), and were thus not included (Supplementary Material, S8).

Lastly, p-values were corrected for multiple-testing, using the Benjamini-Hochberg procedure [48], and their significance was tested against α = 0.05.

## Results

### Quantification of rhythm

Firstly, the density of the interval-ratios was plotted to visualise rhythmic patterning (Figure 1a). The observed interval-ratios of the display behaviour result in a distinct peak at a numerical ratio of 0.5, which marks isochrony. Determined with a Kolmogorov-Smirnov test, the observed ratios differed significantly from the distribution that would have been expected by chance (asymptotic two-sample Kolmogorov-Smirnov test; D = 0.08, *p* < 0.001). Ultimately, this shows that the observed distribution of ratios is not a product of chance.

**Figure 1.**
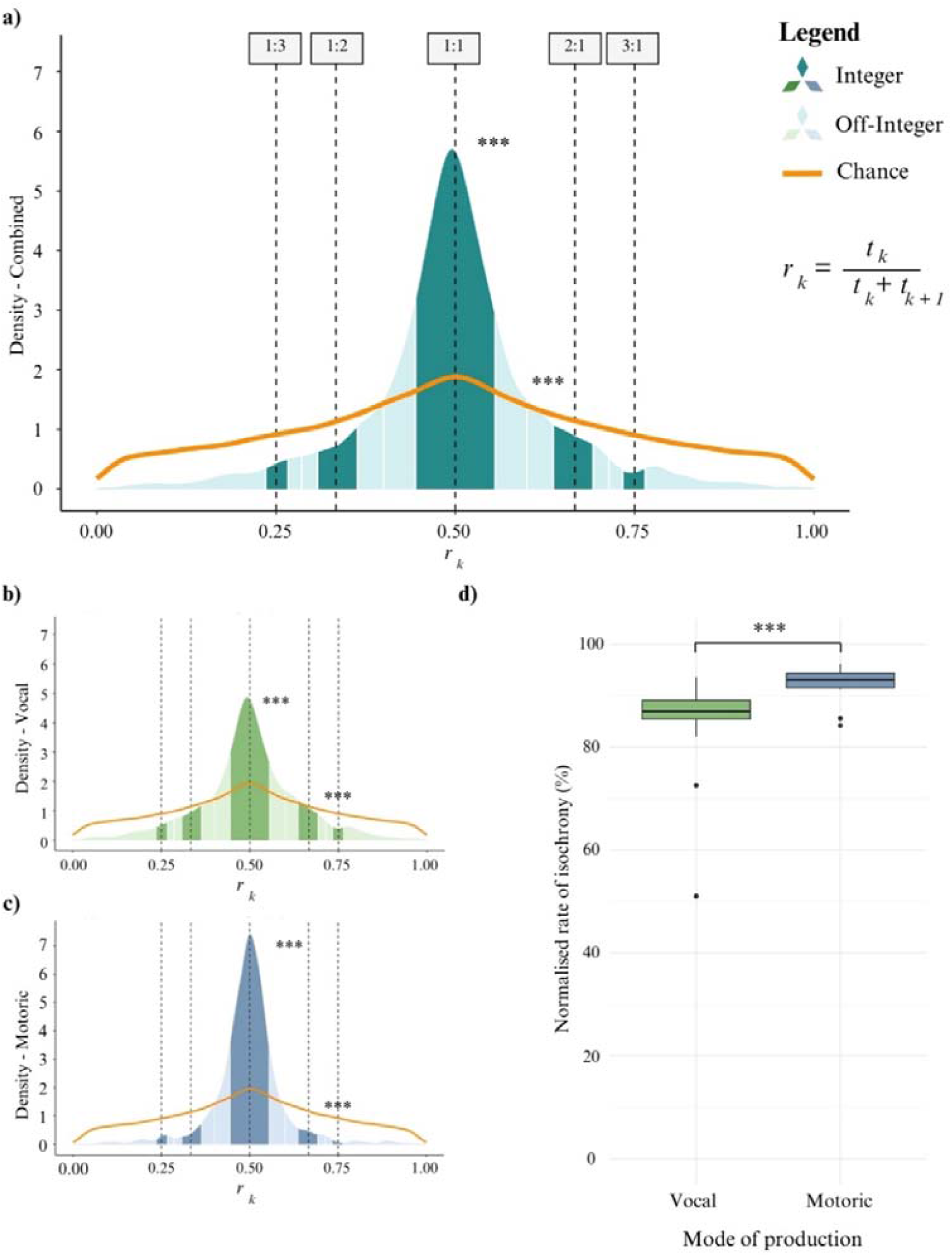
Visualisation of the distribution of the calculated interval-ratios within the observed display behaviour (all behaviours = a, vocal = b, motoric = c). Integer (darker bins) and off-integer (lighter bins) rhythmic categories are visualised with the categorical rhythms 1:3, 1:2, 1:1 (isochrony, rk=0.500), 2:1 and 3:1 marked with dashed lines. More interval-ratios fall within the integer rhythmic boundaries than in the corresponding off-integer bins and the interval-ratios are more likely to be isochronous compared with all other rhythmic categories combined (for a, b & c). The high rates of isochrony in the dataset differ significantly from what would be expected by chance (comparison uniform distribution (orange lines) to ratio-distributions). The motoric elements within the display generally had a higher rate of isochrony than the vocal elements, as depicted in the boxplot with the rate of isochronous interval-ratios (percentage) normalised with bin-size (d).

To compare the normalised counts of ratios in the different categorical rhythms, Wilcoxon signed-rank exact tests were used, as these were not normally distributed (Shapiro-Wilk normality test; W = 0.96, *p* < 0.001). The interval-ratios of the displays fell significantly more within the bounds of isochrony, than in the off-integer bins adjacent to isochrony (Wilcoxon signed-rank exact test; *1:1 ratio*; *integer* versus *off-integer*: V = 78, *p* = 0.001). For the other categorical rhythms, only the interval-ratios within the 1:3 ratio, were significantly higher than the off-integer bins adjacent to this category (Wilcoxon signed-rank exact test; *1:3 ratio*; *integer* versus *off-integer*: V = 72, *p* = 0.012) (Supplementary Material, S5). The number of interval-ratios within isochrony was significantly higher compared to the neighbouring categorical rhythms (Wilcoxon signed-rank exact test; *1:2* versus *1:1*: V = 0, *p* = 0.001; *1:1* versus *2:1*: V = 78, *p* = 0.001). In fact, most of the interval-ratios were concentrated within isochrony, as the count of isochronous ratios was higher than all other off-/integer ratios combined (Wilcoxon signed-rank exact test; *isochrony* versus *off-isochrony*: V = 78, *p* = 0.001).

No significant difference was found between the variability of the inter-onset intervals and the rates of isochrony of the two isolated colonies (LMM, *coefficient of variation; Beekse Bergen versus Burgers’ Zoo:* Estimate =0.048, SE = 0.072, t = 0.665, *p* = 0.516; GLMM, *isochrony rate*; *Beekse Bergen* versus *Burgers’ Zoo*: Estimate = 0.068, SE = 0.237, *Z* = 0.287, *p* = 0.774; Supplementary Material, S4). This suggests that the observed rhythmic patterns remain consistent despite the environmental or genetic differences between the colonies.

### Mode of production

Further, the influence of the mode of production on interval variability (coefficient of variation) and rhythm in the display was evaluated. Interval variability was significantly higher in the vocal sequences than in the motoric sequences (LMM, *coefficient of variation;* motoric versus vocal: Estimate = 0.175, SE = 0.036, *t* = 4.871, p < 0.001; Supplementary Material, S4). Moreover, a significantly higher rate of isochrony was found within the motoric sequences, compared to the vocal sequences (Figure 1b,c: GLMM, *isochrony rate*; *motoric* versus *vocal*: Estimate = -0.892, SE = 0.136, *Z* = -6.537, *p* < 0.001; Supplementary Material, S4). Nevertheless, for both the vocal and motoric sequences, the generated interval-ratios were heavily centred around isochrony. Additionally, comparing both modes to their individually generated chance-distribution, shows that the vocal and motoric ratio density-plots significantly differed from what would be expected by chance (asymptotic two-sample Kolmogorov-Smirnov test; *vocal*: D = 0.08, *p* < 0.001; *motoric*: D = 0.16, *p* < 0.001).

Relatively more interval-ratios were produced in an isochronous rhythm compared to all other categorical rhythms (Wilcoxon signed-rank exact test: *isochrony* versus *off-isochrony*; *vocal*: V = 105, p < 0.001; *motoric*: V = 66, *p* = 0.002). Similarly, the ratio-density was higher within isochrony than in the corresponding off-integer bins (Wilcoxon signed-rank exact test; *1:1 ratio; integer* versus *off-integer*; *vocal*: V = 104, *p* < 0.001; *motoric*: V = 66, *p* = 0.002). In like manner, there was hardly any significant difference between any of the other categorical rhythms and their specific off-integer opposites (Supplementary, S5).

In either mode of production, there were no statistical differences in the ratio-distribution between the different zoos (asymptotic two-sample Kolmogorov-Smirnov test; *Beekse Bergen* versus *Burgers’ Zoo*; *vocal*: D = 0.06, *p* = 0.241; *motoric*: D = 0.08, *p* = 0.319). This demonstrates that the rate of isochrony remains consistent despite variation between/within colonies and in the way in which the display is performed.

### Individually specific tempi

The data indicated variation between the individuals which had at least 25 interval-ratios, vocally (Figure 2a) and/or motorically (Figure 2b). The individuals, from different demographic groups (adult males, adult females and a juvenile male), exhibit varying temporal patterns in their displays (Supplementary Material, S6,7). Specifically, the average duration of the intervals between the elements in a display sequence shows variation within and between individuals, creating individually distinct tempi in the isochronous behaviour. This results in some individuals generally displaying in a faster or slower tempo than other conspecifics. This variation does not seem to be correlated with sex, age, colony or group. Despite the difference in average interval-duration, individuals perform their displays with high rates of isochrony. This indicates that there is no singular tempo at which this rhythmic behaviour is executed, either across different individuals or within the same individual. It also demonstrates that the displays are isochronous, regardless of the displaying individual or the tempo.

**Figure 2.**
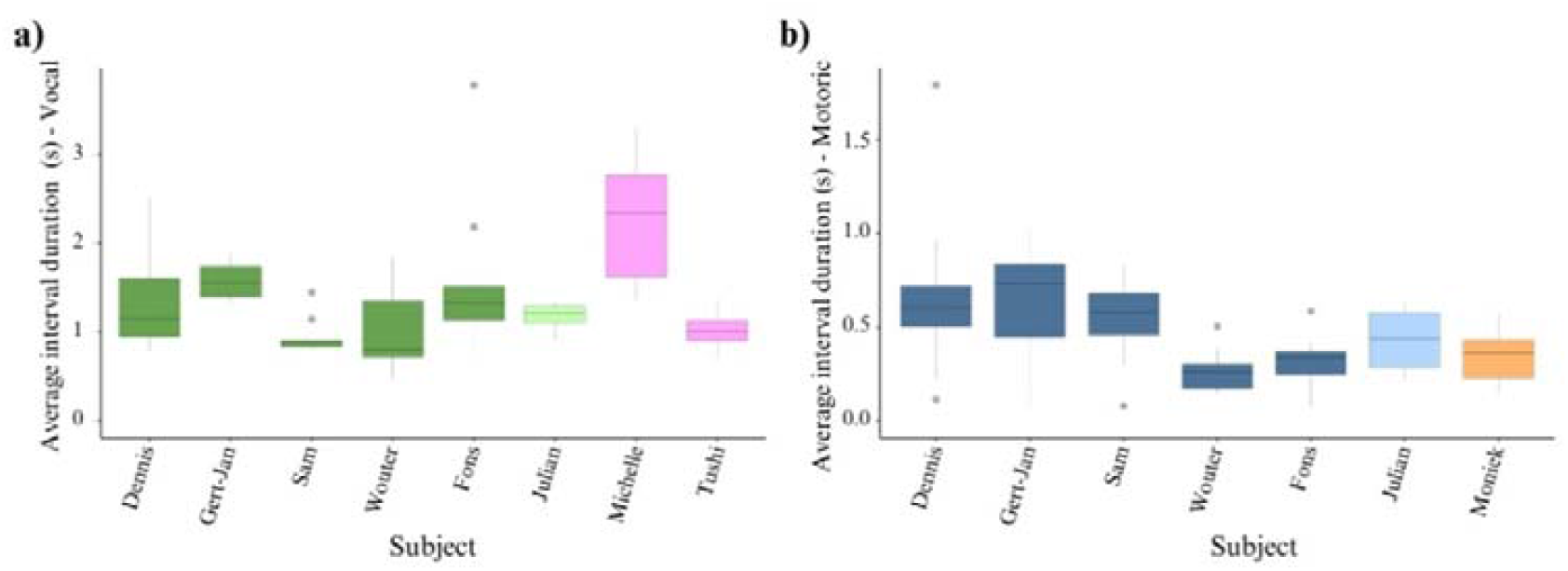
Visualisation of the individual variation in the average duration between intervals (in seconds) per individual for both the vocal (a) and motoric (b) elements of display. This shows the individual differences in tempi of the display behaviour, with different colours that symbolise different demographic groups (adult males, juvenile male, adult females; vocal: dark green, light green, pink; motoric: dark blue, light blue, orange). For the results statistical analysis, the variation in average interval duration between individuals was tested (Supplementary Material, S6,7). Note that the figures have different y-axis scales as the inter-onset-intervals within the motoric displays were, on average, longer.

### Context

Lastly, the influence of the context in which the display behaviour occurred (*directed*: Figure 3a, *undirected*: Figure 3b) and its effect on interval variability and the rate of isochrony, was analysed. Directed and undirected displays did not differ in interval variability (LMM, *coefficient of variation*; *directed* versus *undirected*: Estimate = 0.035, SE = 0.038, *t* = 0.922, *p* = 0.357), but undirected displays were significantly more isochronous than directed displays (Figure 3a and b, GLMM, *isochrony rate*; *directed* versus *undirected*: Estimate = 0.524, SE = 0.153, *Z* = 3.428, *p* < 0.001). This lower rate of isochrony in directed display was notably not correlated with, for instance, lower isochrony rate in vocal displays, as these were mostly undirected (interval-ratios vocal display: *directed* = 503, *undirected* = 1227). Apart from the significant difference in isochrony rate in both contexts, the interval-ratios produced in either context were mainly isochronous.

**Figure 3.**
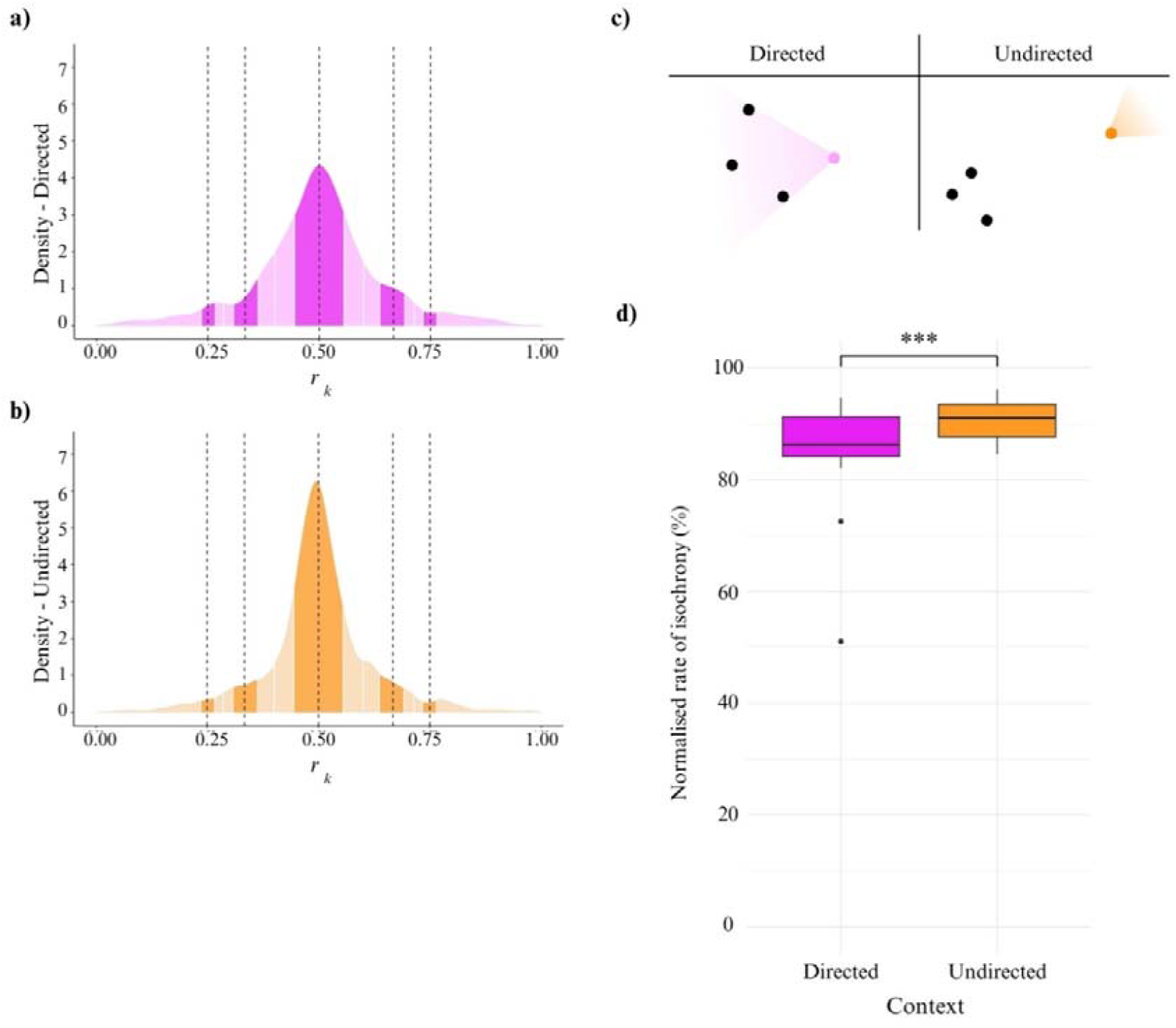
Visualisation of the ratio-distributions of directed displays (a, directed at target individuals (c – left)) and undirected displays (b, oriented towards entire group (c – right)). The bins of the integer categorical rhythms are darker shaded, with the neighbouring off-integer categories being lighter. Both contexts of display show a peak of interval-ratios at isochrony. The undirected displays have a higher rate of isochrony than the displays in a directed context (Supplementary Material, S4), which is depicted in a boxplot of the percentage of isochronous intervals, normalised by the size of the bin of this categorical rhythm (d).

## Discussion

The aim of this study was to quantify the presence and properties of rhythm in display behaviour of zoo-housed chimpanzees and the factors that could influence this rhythm. First and foremost, in the display behaviour, the individuals’ vocal and motoric sequences were both produced in isochronous rhythms. There was clear variation between modes of production, as motoric displays were more regular and had a higher isochrony rate than vocal displays. However, this variation could be (partly) due to the different temporal precision with which the audio recordings of the vocal displays and the video recordings of the motoric displays could be analysed. Furthermore, there was variation in the duration of intervals, resulting in distinct individual display tempi. Nevertheless, overall, the display sequences were highly isochronous.

The clearest variation in rate of isochrony was seen between the two display contexts, directed and undirected, with undirected displays having a higher isochrony rate than directed displays. It should be noted, however, that despite this difference, both contexts showed remarkably high isochrony rates. As undirected displays are not aimed at a target individual, displaying individuals are possibly more focussed on themselves as external stressors are less prominent [32]. Directed displays result inevitably in a divided focus between the action of performing a display and the interaction with targeted individual(s). One could imagine that charging behaviour, following reactive targets during the display, might shift the overall tempo of the behavioural sequence. This might be why isochronous rhythms are maintained relatively less often in directed display sequences. An alternative hypothesis that is proposed in the literature, is that isochrony and regularity in vocalisations or movements have a function in communication. For instance, Australian magpie-larks respond more territorial to regular duets than to irregular duets, regardless of tempo [49]and female mice approach playbacks of regular sequences more than playbacks of irregular sequences [50] However, these findings predict the opposite from what was found in the current study; in the chimpanzees the undirected displays have higher isochrony rates than the directed displays. A study on wild chimpanzees showed that the frequency and structure of pant hoot vocalizations during displays depended on the composition of the audience that these displays were directed to [51]. In the current study all ‘directed’ displays were combined without considering the composition of the audience that these displays were directed to. However, the audience composition might have affected the rhythmic regularity of the displays and future studies should investigate this.

Factors like demography and the specific behaviour performed in the display did not have a significant influence on the rate of isochrony in display sequences. Although adult males did display more often, as was also shown in previous research [27,28], they did not maintain a more consistent rhythm than individuals from other demographic groups. Whilst the influence of these factors cannot fully be ruled out, our findings do suggest that there is a broad foundation for isochronous rhythm production in chimpanzee display behaviour. This rhythm does not seem to be the result of practice, as the youngest individual contributing to the data set was three years old, or even depend on the frequency of displays. Moreover, as individuals uniquely shape their display sequences resulting in a broad behavioural repertoire [27,29–31], the consistent rate of isochrony across individuals in the four observed groups, further signifies the universal base of this rhythmicity.

In the wild, groups of chimpanzees adjust the individual- and group-specific variation in temporal structure of buttress drumming and vocalisations to guarantee inter-/intragroup recognition [22,24,25]. Similarly, implementing variation has been reported in harbour seal pups that tweak their isochronous vocal sequences to produce a distinctly different call in the presence of conspecifics [14]. We found no difference in the regularity or rhythmic rate between the two observed colonies or the different housing-groups within the zoos. This absence of inter-group variation could have been caused by lack of competence for a structural change in rhythm within the entire group. A more plausible explanation, however, is that there was no necessity for groups creating uniquely structured rhythms. The latter is supported by there being hardly any interaction between the two housing-groups in Beekse Bergen. If no interaction occurs between the neighbouring groups, surely it is not essential to differentiate the group-specific rhythmic structure of a display. On top of that, the housing-groups in Burgers’ Zoo were only recently separated (in appreciation of the birth of two young) and rotated between the in- and outside habitat of a single enclosure. Accordingly, individuals from these divided groups likely still recognised each other as members of the same collective group. The interval-tempi between individuals varied similarly to the previously observed interindividual variation in wild chimpanzee behaviour [22,24,25]. This variation confirms that display sequences are flexibly modulated in tempo whilst maintaining a consistently isochronous rhythm, both within and between individuals.

We aimed to quantify rhythmic patterns in both modes of production during chimpanzee display behaviour and found isochronous rhythms despite individual-specific structural variance in behavioural sequences. Since this (or similar) music-based method has been applied in researching rhythmic properties, categorical rhythms were recorded in the vocalisations of a few non-human primates [16,17,19]. We found that chimpanzees share these rhythmic patterns in vocal sequences and that this consistency also manifests itself in the motoric behaviour. These findings provoke additional questions regarding the concept of rhythmicity. Behavioural rhythms can arise from regular physiological oscillations (e.g., heartbeat), be a result of physical constraints (e.g., when a behaviour performed at maximum speed naturally becomes isochronous), or represent the most energy-efficient option. However, some rhythmic behaviours may emerge during higher cognitive processes, where rhythmicity might be generated with a certain intentionality. Although it is complex to differentiate these sources of isochrony, the nature of the behaviour in which rhythmic patterns are exhibited, might offer clues as to whether it stems mainly from physical processes or involves a cognitive component. For example, the difference between the isochrony rate of vocal and motoric behaviours of the chimpanzees, might give some indication to the underlying processes. Assuming that physical restraints have a greater impact on movements than on vocalizations, this could potentially explain why the movements exhibit a more isochronous pattern. However, it should be noted that, as previously mentioned, the rhythmic differences between vocal and motoric behaviour could also be attributed to the different temporal resolution of the analysis of audio and video recordings. Nevertheless, the isochrony in displays is likely an intricate interplay between physical constraints, energy efficiency and potentially an underlying cognitive mechanism.

Rhythmic patterns seem to be present in other social behaviours of chimpanzees and other non-human primates, for instance lip-smacking behaviour that is performed in constant patterns, resembling human speech rhythm [52–55]. Our findings contribute to the expanding documentation that uses music-based methods to quantify rhythmic behaviours across species [12–17,19]. Consequently, the presence of rhythm, unaffected by variation between colonies, individuals, context and mode of production, further underlines the universality of this trait. This is in line with the general hypothesis that rhythm, in human language and music, evolved through a gradual process with simple rhythmic capacities common across species and the advanced skills only present in a few [11]. Future research can investigate these shifts to advanced cognitive capacities, using these music-based methods in a broad range species. This allows for a deeper understanding into how rhythm evolved as the fundamental trait that it is for human language and music.

## Ethics

This study was solely observational, resulting in negligible interference in the daily lives of the recorded chimpanzees. Individuals were only observed during opening hours from, or near, the public route.

## Acknowledgements

First, we want to thank Stijn Berger (Beekse Bergen) for his help and enthusiasm setting up the project. Along with Berger, we want to Kris Jansen (Beekse Bergen), Constanze Mager (Burgers’ Zoo), Safari Park Beekse Bergen and Royal Burgers’ Zoo Arnhem, for facilitating our research by allowing us to observe their chimpanzee populations. This research was supported by ARTIS Amsterdam Royal Zoo, Royal Burgers’ Zoo Arnhem, FWF grant 10.55776/ZK66 & NWO Veni grant 212.264 (both awarded to M.S.).

