## Supplementary material for "Context-dependent Rhythmicity in Chimpanzee Displays"

S1 Group composition, whether subjects displayed and what the contribution was to the sample

| Group | | Subject | Demographic | Year of birth | Performed display? | r_k-_ contribution total | r_k-_ contribution vocal | r_k-_ contribution motoric |
| --- | --- | --- | --- | --- | --- | --- | --- | --- |
| Beekse Bergen | 1 | Dennis  Gert-Jan  Wakili  Sam  Dembe  Mees  Anne-Clara  Linda  Michelle  Marijke  Pepa  Martje  Inocha  Desi | Adult male  Adult male  Adult male  Adult male  Juvenile male  Juvenile male  Adult female  Adult female  Adult female  Adult female  Adult female  Adult female  Adolescent female  Juvenile female | 1984  1995  1999  2003  2020  2021  1980  1985  1996  1997  1999  2000  2014  2022 | Yes  Yes  Yes  Yes  Yes  No  Yes  No  Yes  Yes  No  No  No  No | 87  457  33  540  26  0  17  0  291  27  0  0  0  0 | 44  94  12  495  0  0  13  0  190  25  0  0  0  0 | 43  363  21  45  26  0  4  0  101  2  0  0  0  0 |
|  | 2 | Socrates  Lukani  Daan  Wouter  Julian  Luca  Marlies  Lenny  Nadine  Joke  Lilly  Anna | Adult male  Adult male  Adult male  Adult male  Adolescent male  Juvenile male  Adult female  Adult female  Adult female  Adult female  Adolescent female  Juvenile female | 1981  1988  1997  2000  2009  2021  1981  1982  1990  1996  2015  2020 | Yes  No  Yes  Yes  Yes  Yes    Yes  No  No  No  Yes  Yes | 38  0  36  178  154  17  22  0  0  0  14  12 | 22  0  0  73  106  0  22  0  0  0  0  0 | 16  0  36  105  48  17  0  0  0  0  14  12 |
| Burgers’ Zoo | 1 | Ghineau  Giambo  Roosje  Gaby  Tesua  Morami  Moni  Tushi  Geisha  Laura | Adult male  Adult male  Adult female  Adult female  Adult female  Adult female  Adult female  Adult female  Adult female  Adult female | 2005  1989  1979  1984  1986  1987  1989  1992  1993  1994 | Yes  Yes    Yes  Yes  No  Yes  No  Yes  Yes  Yes | 6  121  3  24  0  1  0  97  4  17 | 2  110  0  0  0  0  0  93  4  17 | 4  11  3  24  0  1  0  4  0  0 |
|  | 2 | Fons  Jing  Moniek  Erika  Raimee | Adult male  Adult male  Adult female  Adult female  Adult female | 1975  1981  1977  1992  1999 | Yes  Yes    Yes  Yes  Yes | 320  42    127  100  180 | 255  35    75  26  169 | 65  7    52  74  11 |

S2 - Table of the characteristics of the study groups of the two zoos, with N representing the animals that cleared the selection-criterion (at least 50 interval-ratios for the total analysis, at least 25 intervals in a specific mode of display production (vocal/motoric) for the analysis of that mode of display production), and were therefore used in the analysis.

|  |  | Total | | After selection criteria |
| --- | --- | --- | --- | --- |
| N | Beekse Bergen  Burgers’ Zoo | 26  15 | | 9  7 |
| Age range  (in years) | Beekse Bergen  Burgers’ Zoo | 1-24  0-48 | | 3-42  24-48 |
| Average age  (in years) | Beekse Bergen  Burgers’ Zoo | 24  36 | | 25  37 |
| N_males_adult_ | Beekse Bergen  Burgers’ Zoo | 8  4 | | 5  3 |
| N_females_adult_ | Beekse Bergen  Burgers’ Zoo | 10  11 | | 2  4 |
| N_males­­_non_adult_ | Beekse Bergen  Burgers’ Zoo | 4  0 | | 2  0 |
| N_females­­_non_adult_ | Beekse Bergen  Burgers’ Zoo | 4  0 | | 0  0 |
| Ratio  contribution | Beekse Bergen  Burgers’ Zoo | 1949  1042 | 1796  987 | |
| Total individuals | | 41 | | 16 |

S3 - Ethogram of the observed behavioural elements for the motoric behaviour within the display. The ‘Element definition’-column states what is seen as a single behavioural element within the sequence of behaviours. The ‘Onset definition’-column states what is seen as the onset of that behavioural element in order to calculate the inter-onset-interval between elements.

| Behaviour | Description | Element definition | Onset definition |
| --- | --- | --- | --- |
| Sway | Swaying, the continuous motion of the subject leaning (mainly the head and shoulders) to either lateral side back and forth, either in a quadrupedal or bipedal stance | The subject has reached the extremity of either lateral side, at which it had started swaying, after completing one full cycle of swaying to the other lateral side and back | The moment at which the subjects switches the lateral direction of the swaying motion and starts swaying towards the other lateral side |
| Bounce | Bouncing, the continuous motion of the subject moving (mainly the head and shoulders) up and down, either in a quadrupedal or bipedal stance | The subject has reached the most up- or downward position, at which it had started moving, after completing a full cycle of bouncing up- or down to the other side and back | The moment at which the subject switches the direction of bouncing motion and starts moving towards to the other side, either up or down |
| Rock | Rocking, the continuous motion of the subject leaning (mainly the head and shoulders) forwards and backwards, either in a quadrupedel or bipedal stance | The subject has reached the most forward- or backwards position, at which it had started leaning, after completing a full cycle of rocking forwards and backwards | The moment at which the subject switches the direction of the rocking motion and starts moving towards to other side, either forwards or backwards |
| Stomp Right Foot | Hitting the ground or the surface on which the animal is standing, hard with the right foot | A single instance of the subject hitting the ground or surface on which the animal is standing, hard with its right foot | The moment at which the right foot of the animal makes contact with the ground or floor surface |
| Stomp Left Foot | Hitting the ground or the surface on which the animal is standing, hard with the left foot | A single instance of the subject hitting the ground or surface on which the animal is standing, hard with its left foot | The moment at which the left foot of the animal makes contact with the ground or floor surface |
| Stomp Right Hand | Hitting the ground or the surface on which the animal is standing, hard with the right hand | A single instance of the subject hitting the ground or surface on which the animal is standing, hard with its right hand | The moment at which the right hand of the animal makes contact with the ground or floor surface |
| Stomp Left Hand | Hitting the ground or the surface on which the animal is standing, hard with the left hand | A single instance of the subject hitting the ground or surface on which the animal is standing, hard with its left hand | The moment at which the left hand of the animal makes contact with the ground or floor surface |
| Hit Wall Right Foot | Kicking a vertical surface,e.g. a wall, door or plank, with the right foot | A single instance of the subject hitting the vertical surface, e.g. a wall, door or plank, hard with its right foot | The moment at which the right foot makes contact with the wall, door or plank |
| Hit Wall Left Foot | Kicking a vertical surface, e.g. a wall, door or plank, with the left foot | A single instance of the subject hitting the vertical surface, e.g. a wall, door or plank, hard with its left foot | The moment at which the left foot makes contact with the wall, door or plank |
| Hit Wall Right Hand | Hitting a vertical surface, e.g. a wall, door or plank, with the right hand | A single instance of the subject hitting the vertical surface, e.g. a wall, door or plank, hard with its right hand | The moment at which the right hand makes contact with the wall, door or plank |
| Hit Wall Left Hand | Hitting a vertical surface, e.g. a wall, door or plank, with the left hand | A single instance of the subject hitting the vertical surface, e.g. a wall, door or plank, hard with its left hand | The moment at which the left hand makes contact with the wall, door or plank |
| Hit Wall Not Visible | Hitting a vertical surface, e.g. a wall, door or plank, hard with either an indeterminable limb or a limb out of frame | A single instance of the subject audibly hitting the vertical surface, e.g. a wall, door or plank with an indeterminable limb or a limb out of frame. | The moment at which the sound of impact with the wall is audible |
| Clap | Clapping, striking the palms of the hands together, possibly creating a sound | A single clap: a single instance of the palms of the hands forcefully making contact with one another | The moment at which the palms of the hand make contact with one another |
| Kick Right Leg | Kicking the right leg in the airin a hitting motion, without making physical contact with another individual, object or surface | A single instance of moving the right leg in a kicking motion, as if trying to hit another individual, object or surface, without making physical contact | The moment at which the subject had reached the extreme point of extension of the right leg, without making physical contact, before retracting the right leg again |
| Kick Left Leg | Kicking the left leg in the airin a hitting motion, without making physical contact with another individual, object or surface | A single instance of moving the left leg in a hitting motion, as if trying to hit another individual, object or surface, without making physical contact | The moment at which the subject had reached the extreme point of extension of the left leg, without making physical contact, before retracting the left leg again |
| Swing Right Arm | Swinging the right arm in the air in a hitting motion, without making physical contact with another individual, object or surface | A single instance of moving the right arm in a hitting motion, as if trying to hit another individual, object or surface, without making physical contact | The moment at which the subject had reached the extreme point of extension of the right arm, without making physical contact, before retracting the right arm again |
| Swing Left Arm | Swinging the left arm in the air in a hitting motion, without making physical contact with another individual, object or surface | A single instance of moving the left arm in a hitting motion, as if trying to hit another individual, object or surface, without making physical contact | The moment at which the subject had reached the extreme point of extension of the left arm, without making physical contact, before retracting the left arm again |
| Slap Right Hand | Striking another individual with the right hand and making physical contact in this motion | A single instance of hitting another individual with the right hand | The moment at which the subject makes contact with the targetted individual with the right hand in a hitting motion |
| Slap Left Hand | Striking another individual with the left hand and making physical contact in this motion | A single instance of hitting another individual with the left hand | The moment at which the subject makes contact with the targetted individual with the left hand in a hitting motion |
| Grab Object | Grabbing an object, e.g. a stick, branch or piece of plastic | A single instance of the subject grabbing an object, until the object has fully been let go | The moment at which the subject makes contact with the object in order to grab it |
| Pull Object | The motion of pulling on an object, e.g. a branch, by shifting the bodily weight or retracting the limb which is holding the object | A single instance of the subject exerting force on the object by e.g. retracting the limb which is holding the object, until a continuous motion of pulling on the object is stopped | The moment at which the subject starts the motion that exerts force on the object, e.g. retracting the arm |
| Wave Object Right Arm | Waiving an object with the right arm, the continuous motion of alternating waiving movement with right arm, whilst holding an object, e.g. a branch | The subject has reached the extremity of either lateral side, or forwards or backwards, at which it had started waving its right arm, whilst holding an object, after completing one full cycle of waving the held object towards the other side and back | The moment at which the subject has reached the extreme end in the axis in which it is moving its right arm, whilst holding an object in that arm |
| Wave Object Left Arm | Waiving an object with the left arm, the continuous motion of alternating waiving movement with left arm, whilst holding an object, e.g. a branch | The subject has reached the extremity of either lateral side, or forwards or backwards, at which it had started waving its right arm, whilst holding an object, after completing one full cycle of waving the held object towards the other side and back | The moment at which the subject has reached the extreme end in the axis in which it is moving its left arm, whilst holding an object in that arm |
| Throw Object | Throwing an object, which was previously held, e.g. a stick or a piece of plastic, undirectedly | A single instance of the subject letting go of the object whilst forcefully moving its arm, throwing the object away | The moment at which there is no longer any contact between the object and the subject |
| Throw Object Directed | Throwing an object, which was previously held, e.g. a stick or a piece of plastic, at another individual | A single instance of the subject letting go of the object whilst forcefully moving its arm, throwing the object towards another individual | The moment at which there is no longer any contact between the object and the subject |
| Raise Right Arm | The subject lifts its right arm and raises it above its head and shoulders | The instance of the subject raising its right arm above its head, until the arm is lowered below this range again | The moment at which the subject starts the motion of moving its right arm upwards |
| Raise Left Arm | The subject lifts its left arm and raises it above its head and shoulders | The instance of the subject raising its left arm above its head, until the arm is lowered below this range again | The moment at which the subject starts the motion of moving its left arm upwards |
| Wave Right Arm | Waving right arm, continuous motion of alternating lateral movement of the raised right arm | The subject has reached the extremity of either lateral side with its raised right arm, at which it had started waving, after completing one full cycle of waving the right arm to the other lateral side and back | The moment at which the subjects switches the lateral direction of the waving motion with its raised right arm and starts waving the right arm towards the other lateral side |
| Wave Left Arm | Waving left arm, continuous motion of alternating lateral movement of the raised left arm | The subject has reached the extremity of either lateral side with its raised left arm, at which it had started waving, after completing one full cycle of waving the right arm to the other lateral side and back | The moment at which the subjects switches the lateral direction of the waving motion with its raised left arm and starts waving the left arm towards the other lateral side |

S4 Output of statistical models to analyse the effect of different factors on interval variability and isochrony rate in the displays. This was analysed in R (version 4.2.2) [1] with R packages lme4 [2]; function lmer [3]; Model assessment; R packages performance [4], DHARMa [5]. The significant correlations are highlighted in grey.

*a LMM output of the fixed effects of the associations between the unbiased coefficient of variation for the intervals in the behavioral sequences and the form production, directedness of the display and the zoo where the behaviour was observed, with the ID of the displaying individual as random effects.*

| Groups | Estimate | SE | *t* | *p* |
| --- | --- | --- | --- | --- |
| (Intercept) | 0.330 | 0.051 | 6.475 | < 0.001 |
| Form of Production (Vocal) | 0.175 | 0.036 | 4.871 | < 0.001 |
| Directedness (Undirected) | 0.035 | 0.038 | 0.922 | 0.357 |
| Zoo (Burgers’ Zoo) | 0.048 | 0.072 | 0.665 | 0.516 |

b GLMM output of the fixed effects of the associations between isochrony in the display (set as 1 in binomial link function) and the form production, directedness of the display and the zoo where the behaviour was observed, with the ID of the display and the displaying individual as random effects. This was analysed in R (version 4.2.2) [1] with R packages lme4 [2]; function lmer [3]; Model assessment; R packages performance [4], DHARMa [5]. The significant correlations are highlighted in grey.

| Groups | Estimate | SE | *Z* | *p* |
| --- | --- | --- | --- | --- |
| (Intercept) | 0.279 | 0.190 | 1.465 | 0.143 |
| Form of Production (Vocal) | -0.892 | 0.136 | -6.537 | < 0.001 |
| Directedness (Undirected) | 0.524 | 0.153 | 3.428 | < 0.001 |
| Zoo (Burgers’ Zoo) | 0.068 | 0.237 | 0.287 | 0.774 |

S5 - Output of the statistical analysis of the interval-ratio distribution within the categorical rhythms. The normalised count of interval-ratios within a categorical rhythm was compared against the count in another category, in order to analyse the distribution of the interval-ratios. This analysis was performed via multiple Wilcoxon signed rank exact tests and corrected for multiple testing via the Benjamini-Hochberg procedure [6]. The significant correlations have been highlighted in grey. Significant correlations regarding isochrony, the ratio of 1:1 and the different categorical rhythms compared to each other assert the isochronous peak of the distribution of the interval-ratios. The significant difference between the 1:3 and 1:2 ratios combined (slowing down) compared to the 2:1 and 3:1 ratios combined (speeding up) in the motoric elements of display suggests that the subjects generally slowdown in interval speed whilst displaying motorically more than they speed up. This was not mentioned as this result does not uphold in the other categories of display (in vocalisations and in both modes of production combined), as a matter of fact, here, this correlation is strongly non-significant. This correlation might be a result of motoric fatigue whilst displaying. A more plausible explanation, however, is that this effect is likely the result of the stronger peak at isochrony in the motoric mode of display production, compared to the other distributions. This would quickly result in a bigger difference between the other categorical rhythms, as there are relatively fewer interval-ratios within these categories.

| Multiple Wilcoxon signed rank exact tests (B-H correction) | Total | | Vocal | | Motoric | |
| --- | --- | --- | --- | --- | --- | --- |
| *Isochrony* versus  *Off-Isochrony* | V = 78 | *p* = 0.001 | V = 105 | *p* < 0.001 | V = 66 | *p* = 0.002 |
| *Ratio 1:1* *Integer* versus *Off-Integer* | V = 78 | *p* = 0.001 | V = 104 | *p* < 0.001 | V = 66 | *p* = 0.002 |
| *Ratio 1:3 Integer* versus *Off-Integer* | V = 72 | *p* = 0.012 | V = 82 | *p* = 0.018 | V = 30 | *p* = 0.446 |
| *Ratio 1:2 Integer* versus *Off-Integer* | V = 13 | *p* = 0.061 | V = 34 | *p* = 0.319 | V = 14 | *p* = 0.235 |
| *Ratio 2:1 Integer* versus *Off-Integer* | V = 21 | *p* = 0.226 | V = 47 | *p* = 0.804 | V = 14 | *p* = 0.383 |
| *Ratio 3:1 Integer* versus *Off-Integer* | V = 52 | *p* = 0.383 | V = 65 | *p* = 0.235 | V = 20 | *p* = 0.848 |
| *Ratio 1:3* versus *Ratio 1:2* | V = 12 | *p* = 0.050 | V = 8 | *p* = 0.016 | V = 21 | *p* = 0.580 |
| *Ratio 1:2* versus *Ratio 1:1* | V = 0 | *p* = 0.001 | V = 1 | *p* < 0.001 | V = 0 | *p* = 0.007 |
| *Ratio 1:1* versus *Ratio 2:1* | V = 78 | *p* = 0.001 | V = 104 | *p* < 0.001 | V = 66 | *p* = 0.007 |
| *Ratio 2:1* versus *Ratio 3:1* | V = 78 | *p* = 0.001 | V = 102 | *p =* 0.004 | V = 19 | *p* = 0.128 |
| *Slowing down  (Ratio 1:3 & 1:2 combined)* versus *Speeding up (Ratio 2:1 & 3:1 combined)* | V = 562 | *p* = 0.333 | V = 645 | *p* = 0.946 | V = 291 | *p* = 0.022 |

S6 - The p-values of pairwise Kruskal-Wallis rank sum test (Chi-squared = 29.49, df = 7, p < 0.001), performing an inter-individual comparison of the average interval duration per display of the vocalisations within the display behaviour. After the subject name, the number of recorded instances of display behaviour from this individual has been stated (N). The p-values shown have been adjusted for multiple testing with Benjamini-Hochberg procedure. The significant correlations are highlighted in grey.

| Subject (N) | Dennis (4) | Fons (13) | Gert-Jan (4) | Julian (7) | Michelle (11) | Sam (10) | Tushi (4) |
| --- | --- | --- | --- | --- | --- | --- | --- |
| Fons (13) | 0.646 |  |  |  |  |  |  |
| Gert-Jan (4) | 0.486 | 0.495 |  |  |  |  |  |
| Julian (7) | 0.925 | 0.332 | 0.057 |  |  |  |  |
| Michelle (11) | 0.120 | 0.041 | 0.221 | 0.008 |  |  |  |
| Sam (10) | 0.476 | 0.041 | 0.047 | 0.047 | 0.005 |  |  |
| Tushi (4) | 0.716 | 0.221 | 0.154 | 0.657 | 0.042 | 0.658 |  |
| Wouter (5) | 0.658 | 0.350 | 0.332 | 0.701 | 0.043 | 0.679 | 0.925 |

S7 - The p-values of pairwise Kruskal-Wallis rank sum test (Chi-squared = 28.18, df = 6, p < 0.001), performing an inter-individual comparison of the average interval duration per display of the motoric behaviour within the display behaviour. After the subject name, the number of recorded instances of display behaviour from this individual has been stated (N). The p-values shown have been adjusted for multiple testing with Benjamini-Hochberg procedure. The significant correlations are highlighted in grey.

| Subject (N) | Dennis (10) | Fons (12) | Gert-Jan (22) | Julian (12) | Moniek (6) | Sam (11) |
| --- | --- | --- | --- | --- | --- | --- |
| Fons (12) | 0.040 |  |  |  |  |  |
| Gert-Jan (22) | 0.500 | 0.008 |  |  |  |  |
| Julian (12) | 0.160 | 0.151 | 0.040 |  |  |  |
| Moniek (6) | 0.121 | 0.761 | 0.040 | 0.348 |  |  |
| Sam (11) | 0.880 | 0.040 | 0.209 | 0.207 | 0.160 |  |
| Wouter (11) | 0.040 | 0.348 | 0.008 | 0.054 | 0.416 | 0.040 |

S8 - GLMM output of unused model for the fixed effects of the associations between isochrony in the display (set as 1 in binomial link function) and the form production, directedness of the display, the zoo where the behaviour was observed, group in which the animals were housed and the demographic identity of the animals, with the ID of the display and the displaying individual as random effects. This model was ultimately not used in the final analysis, since the addition of the fixed effects of the housing groups and the demographic identity of the animals did not improve the fit of the model. The factor of the type of behaviour, i.e. call type and the type of motoric element, were not included as these did not only decrease the fit of the model, but were also highly multicollinear.

| GLMM | Estimate | | SE | | *Z* | | *p* |
| --- | --- | --- | --- | --- | --- | --- | --- |
| (Intercept) | 0.376 | 0.270 | | 1.394 | | 0.163 | |
| Form of Production (Vocal) | -0.904 | 0.138 | | -6.530 | | < 0.001 | |
| Directedness (Undirected) | 0.524 | 0.159 | | 3.306 | | < 0.001 | |
| Zoo (Burgers’ Zoo) | -0.068 | 0.279 | | -0.245 | | 0.806 | |
| Group (Group 2) | -0.059 | 0.392 | | -0.149 | | 0.881 | |
| Group (Group A) | 0.220 | 0.361 | | 0.609 | | 0.543 | |
| Demographic (Adult Male) | -0.056 | 0.256 | | -0.219 | | 0.827 | |
| Demographic (Juvenile Male) | -0.199 | 0.530 | | -0.376 | | 0.707 | |
